# Using the ancestral recombination graph to study the history of rare variants in founder populations

**DOI:** 10.1101/2025.03.13.643149

**Authors:** Alejandro Mejia Garcia, Alex Diaz-Papkovich, Guillaume Sillon, Daniela D’Agostino, Anne-Laure Chong, George Chong, Ken Sin Lo, Laurence Baret, Nancy Hamel, Vincent Chapdelaine, William D. Foulkes, Daniel Taliun, Adam J. Shapiro, Guillaume Lettre, Simon Gravel

## Abstract

Gene genealogies represent the ancestry of a sample and are often encoded as ancestral recombination graphs (ARG). It has recently become possible to infer these gene genealogies from sequencing or genotyping data and use them for evolutionary and statistical genetics. Unfortunately, inferred gene genealogies can be noisy and subject to biases, making their applications more challenging. This project aims to study the application of ARG methods to systematically impute and trace the transmission of all disease variants in founder populations where long-shared haplotypes allow for accurate timing of relatedness. We applied these methods to the population of Quebec, where multiple founder events led to an uneven distribution of pathogenic variants across regions and where extensive population pedigrees are available. We validated our approach with nine founder mutations for the SLSJ region, demonstrating high accuracy for mutation age, imputation, and regional frequency estimation. Moreover, we showed that this subset of high-quality carriers is sufficient to capture previously described associations with pathogenic variants in the *LPL* gene. This method systematically characterizes rare variants in founder populations, establishing a fast and accurate approach to inform genetic screening programs.

## Introduction

A disease is considered rare if it affects fewer than 200.000 people (or 1 in 1.650) in the United States (1) and fewer than 1 in 2.000 people in Europe (2). In aggregate, 7.000 rare diseases affect 7% of the human population (1). Around 80% of rare diseases are thought to be monogenic (3). Alleles causing rare diseases tend to have a low frequency in the population (<1%) (4). These alleles can be difficult to study in population cohorts due to limited statistical power and phenotype heterogeneity(4,5).

For cohorts using genotyping array data, the problem is compounded by challenges in imputation of rare variants, even when the variant is present in reference panels (6). The presence of rare variants at higher frequency in a founder population can allow for improved association power (7). Because multiple carriers may share a recent common ancestor, the shared haplotype structure can lend itself to improved imputation, as well as evolutionary studies (8). Where genealogical data is available, detailed transmission patterns can sometimes be inferred (9).

The Ancestral Recombination Graph, or ARG, is a unified representation of the shared haplotype structure of all variants across the genome (10,11). Recent work has made it possible to infer the ARG at biobank scale, including from ascertained genotype data. Inferred ARGs have been used in association studies, demographic inference, evolutionary studies, and data compression, among others (10,11).

The goal of this article is to investigate the use of inferred ARGs for the study of rare variants in a founder population, including imputation, association, and inference of transmission histories within documented genealogies. We will focus on the population of Quebec, Canada. The territory of present-day Quebec has been inhabited by Indigenous communities for thousands of years (12–15). The migration and eventual settlement of around 8.500 individuals, mostly from western and northwestern France, primarily to Quebec City (1608) and Montréal (1642) (15,16), led to a series of founder events (12,16). After the British conquest in 1760, French migration ceased, and the French-Canadian population expanded rapidly with relative isolation through linguistic and religious barriers (14,15,17). This population growth led to the colonization of new regions, favouring population subdivisions (12,18). For example, migrations to Saguenay-Lac-Saint-Jean (SLSJ) primarily from the Charlevoix region, led to another well-known regional founder event (17).

These demographic events are explicitly documented by extensive genealogical records through BALSAC, a genealogy reconstructed from birth, death, and marriage records, with a depth of up to 17 generations, and includes most early French settlers (19).

Because of regional founder events, the frequency of rare variants varies across Quebec (12,20,21). In SLSJ, Charlevoix and Haute-Côte-Nord regions, for example, a screening program exists to test for variants causing hereditary tyrosinemia type I, autosomal recessive spastic ataxia of Charlevoix-Saguenay (ARSACS), Leigh-syndrome French-Canadian type (LSFC) and agenesis of corpus callosum and peripheral neuropathy (ACCPN) (22,23).

However, several founder pathogenic variants causing other diseases like cancer, neuropathy, glaucoma, Tay-Sachs disease and Usher syndrome, among others, have been reported in Quebec, but details available about their transmission history and their frequency across the province are not always well documented (21,24–28).

Recently, the CARTaGENE cohort generated genotype data for 29,337 individuals of diverse ancestries (29) and performed high-depth whole-genome sequencing (WGS) of a subset of over 2.173 individuals. Moreover, 7.894 of these individuals are linked to the BALSAC dataset (19).

In this study, we generated the ARG for the 30,000 genotyped individuals. We show how this inferred ARG can be used to assist in the systematic analysis of variants of interest at frequencies above 1/500 in the cohort, including for imputation, association, and estimation of regional allele frequencies. Concretely, many of our analyses use the ARG to identify haplotypes likely introduced in the population by a single individual, enabling straightforward imputation and transmission inference using tools such as ISGEN (9).

## Subjects and Methods

### CARTaGENE cohort

The CARTaGENE (CaG) longitudinal cohort recruited 43.000 participants (40-69 years old at recruitment) across five different regions in the Province of Quebec, Canada. The cohort has been described elsewhere (29). 29.337 individuals have genotype data available from different GSA arrays (https://cartagene.qc.ca/files/documents/other/Info_GeneticData3juillet2023.pdf) that were unified in a single file with PLINK 1.9 (30), and, after quality control, a total of 659,029 variants were included. Data was then imputed using the TOPMed Imputation Server (panel TOPMed-r2, n=97,256 deeply sequenced human genomes), resulting in a total of 109,242,846 variants.

In addition, a subset of individuals (n=2,173) has high-depth WGS data available, including 1,756 individuals with four grandparents born in Canada with French as a primary language, as well as 163 and 132 participants with four grandparents born in Haiti and Morocco, respectively.

Moreover, extensive questionnaire data and blood measurements, including white blood cells, lipid profile, glucose, among others, were available for 29,143 individuals (29).

### BALSAC genealogy

The BALSAC genealogical database was constructed mainly from birth, death, and marriage certificates and contains around 6.5M individuals from the 1600s to the present (19). The maximum depth of the genealogy is 17 generations. Most of the ancestry of present-day Quebec residents can be traced back at least 12 generations (19) and the reconstruction is particularly complete among Quebec residents of French-Canadian ancestry (QFC). Moreover, 7,894 individuals from CARTaGENE are linked to this genealogy.

### ARG reconstruction

To take advantage of the extensive quality control performed previously, we used imputed VCF files (109,242,846 variants) from 29,337 individuals and filtered them to include only genotyped variants with phase information using bcftools (727,936 variants) (31). We excluded chromosomes X and Y from the analysis, leaving 659,029 variants. We then kept only biallelic SNPs using plink2 (30) –max-alleles 2 and --snps-only flags, leaving 566,073 variants. Moreover, we used genetic map files for GRCh38 reference genome to obtain the map position of all SNPs using an in-house R script (https://github.com/almejiaga/ARG_needle). We discarded variants with a distance lower than 0.00001cM due to ARG-needle software requirements. Finally, we converted VCF files to Oxford HAP format using plink2 (30). In the end, 548,663 variants were retained for subsequent analysis. We reconstructed the ARG using the ARG-needle software (10) in genotype mode with default parameters for chromosomes 1-22. We converted ARG files tree sequences in tskit format using the arg2tskit function of ARG-needle.

The 2,173 individuals with WGS data also had genotype array data available. For consistency, the ARG was reconstructed using only genotype data.

### ARG-based imputation

For any variant x, we first identified carriers in the WGS data, and we computed the most recent common ancestor node (MRCA) between them in the ARG using the tree.mrca tskit function (https://github.com/tskit-dev/tskit/). Given statistical phasing uncertainty for rare variants, we used an ARG-based heuristic phasing strategy: given a number of carriers of a rare variant, it is very likely that at least a few carrier haplotypes share a recent MRCA. By contrast, the non-carrier haplotypes are unlikely to all share a recent MRCA. Thus, for a group of K diploid carriers (with 2K haplotypes), we select a subclade of K haplotypes with the lowest ARG-inferred time to the most recent common ancestor (TMRCA, labelled as *t*_1_ on Figure 1a), we labelled the corresponding node as the carrier MRCA (cMRCA). We then selected the cMRCA as a likely carrier. We assume that the mutation occurred along a branch ancestral to the cMRCA. We also assume that it occurred below the common ancestor of the cMRCA with sequenced non-carriers, at time *t*2(Figures 1a and b).

**Figure 1.**
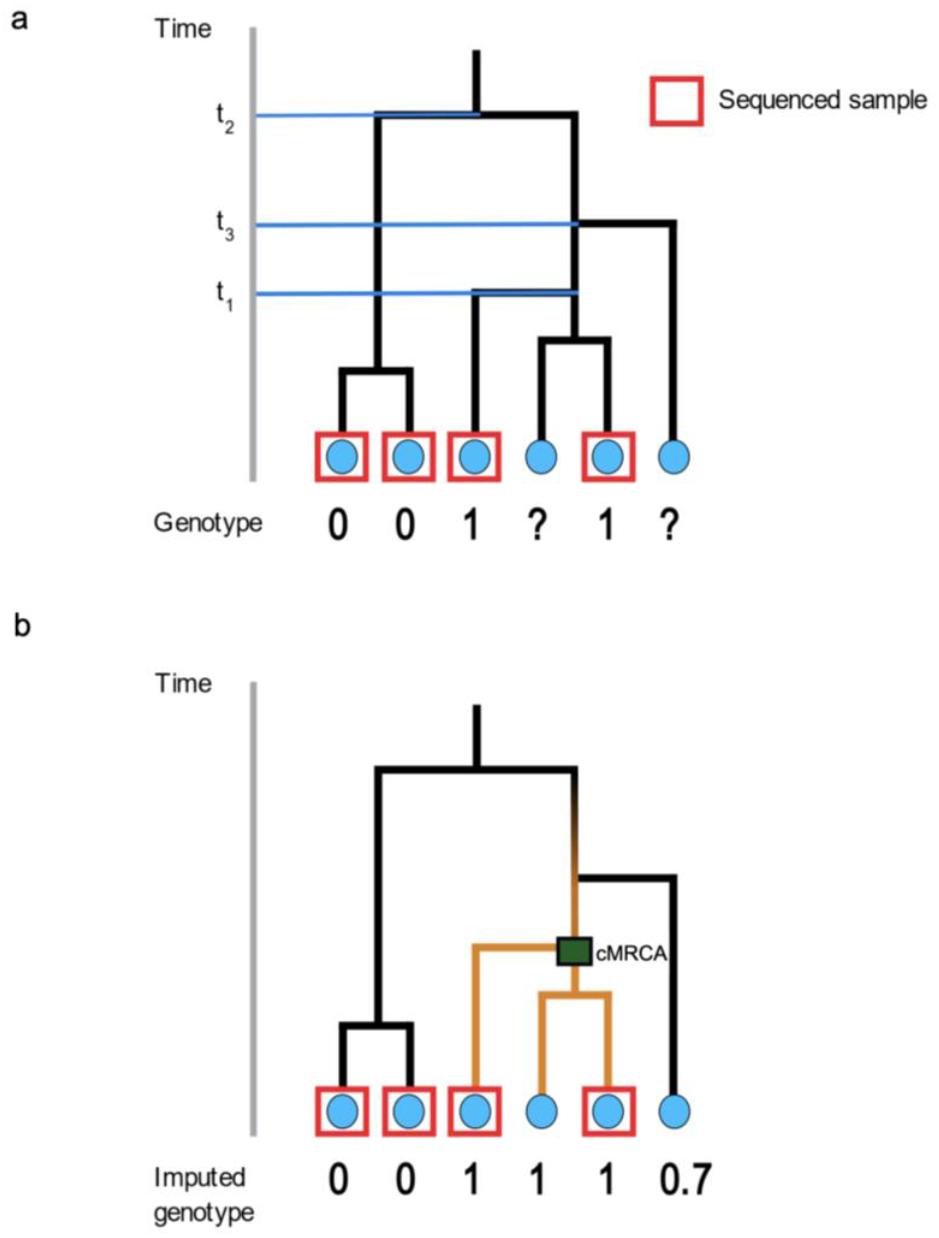
Overview of the ARG-imputation method. a and b) We find the MRCA between the set of carriers (green). We assume that all individuals below the MRCA carry the mutation, and they become imputed carriers P (c) = 1. Individuals with a TMRCA between t1 and t2 get a posterior probability with Equation 1. For individuals with a TMRCA >= t_2_, P (c) = 0.

For imputation, we suppose that all the descendants of the cMRCA carry the variant. For individuals connecting to the ARG above the cMRCA at *t*_3_ ≤ *t*_2_, we estimated the posterior probability of being a carrier *C* (t_3_, Figure 1c) using the following equation:

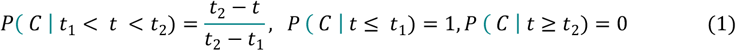

### Clade mixing score

In this procedure, there may be sequenced individuals who are not carriers but are descendants to the cMRCA. If *S* is the set of descendant nodes to the cMRCA at a locus, we define the mixing score

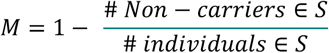

Since these non-carriers would have been imputed as carriers, *M* can be thought of as an estimate of imputation false positive rate.

### Experimental validation of ARG-imputation

To independently validate our imputation strategy, we applied ARG-imputation to the *PMS2 c*.*2117del* and *ATM c*.*7388_7389insAlu p*.*Leu2463fs* pathogenic variants in the whole CARTaGENE cohort (n=29,337). To assess its accuracy, we directly genotyped a subset of 4,000 individuals from CARTaGENE for these two pathogenic variants by Sanger sequencing, using this genotyping data as the gold standard. We then compared the ARG-imputation results with the Sanger-genotyped data, evaluating imputation performance using accuracy and the Kappa statistic.

### Comparison of ARG-imputation against imputation with a Quebec reference panel

To further validate our approach, we compared against a standard imputation method (Minimac4). WGS individuals (n=2,172) were used as reference for imputation using Minimac4 (32). We then compared carriers assigned by this imputation against ARG-imputation. We built contingency tables and computed the following agreement metrics: accuracy and Kappa statistic. In addition, we visualized the posterior probability distribution computed from the ARG for Minimac4 carriers.

### Genotype-phenotype association analysis with *LPL: c*.*701C>T* carriers

We performed an association between the carrier posterior probability (Equation 1) and triglycerides, high-density lipoprotein cholesterol (HDL-C), and Low-density lipoprotein cholesterol (LDL-C) levels, among other blood lipid measurements using linear regression with age and sex as covariates, as implemented in statsmodels python package. P-values were adjusted using Bonferroni correction to account for multiple testing, assuming independent tests.

### The carrier frequency in Saguenay–Lac-Saint-Jean

Many regional founder variants have been documented in the Saguenay-Lac-Saint-Jean region of Quebec. To demonstrate that imputed carriers could be used to estimate allele frequencies in Saguenay, we estimated the carrier frequency for nine known founder pathogenic variants in Saguenay, four of which are included in the carrier screening program in Quebec (22,23). We used genetic clustering to identify individuals with likely ancestry in SLSJ. We applied UMAP using the top 10 PCs from principal component analysis (PCA) and 15 nearest neighbors and a minimum distance of 0.5. We then clustered using HDBSCAN with the following parameters: *ε*=0.3, minimum cluster size of 25 on a 3D UMAP of the top 25 PCs parameterized for the 10 nearest neighbours and a minimum distance of 0.001, as described in (33). We used clusters defined by HDBSCAN to estimate carrier frequency in SLSJ in two different types of data: WGS and ARG-imputed genotypes. We compared their frequency against the literature and ISGen estimations (Table 1), explained below.

**Table 1:** Inherited disorders in Saguenay–Lac-Saint-Jean (SLSJ)

| Autosomal Recessive disorders |  |  |  |  |  |  |  |  |  |
| --- | --- | --- | --- | --- | --- | --- | --- | --- | --- |
| Disease | Variant | TMRCAs | GnomAD NFE | CARTaGENE |  | Saguenay carrier frequency |  |  |  |
|  |  |  |  | WGS | Imputation | WGS | ARG imputation | ISGen | Literature (21) |
| Mucopolipidosis type 2 | GNPTAB: c.3503_3504delTC | 26.94 | 1/927 | 8 | 84 | 1/42 | 1/51 | 1/31 | 1/39 |
| Zellweger syndrome | PEX6c.802-825del14 | 14.72 | 1/49168 | 4 | 56 | 1/62.5 | 1/83 | 1/80 | 1/55 |
| Tyrosinemia type I | FAH:c.1062+5G>A | 31.42 | 1/1077 | 19 | 219 | 1/15 | 1/15 | 1/16 | 1/20 |
| Cystinosis | CTNS:c.414G>A (p.Trp138X) | 16.48 | 1/5413 | 7 | 94 | 1/42 | 1/45 | 1/36 | 1/39 |
| Vitamin D-dependent rickets Type I | CYP27B1:c.262del (p.Val88fs) | 16.61 | 1/12766 | 11 | 202 | 1/28 | 1/26 | 1/33 | 1/27 |
| ARSACS* | SACS:c.8844del (p.Ile2949fs) | 13.25 | 1/13404 | 24 | 277 | 1/27 | 1/18 | 1/16 | 1/22 |
| LSFC** | LRPPRC:c.1061C>T (p.Ala354Val) | 6.03 | 1/17306 | 7 | 119 | 1/125 | 1/33 | 1/45 | 1/23 |
| ACCPN*** | SLC12A6:c.2436delG (p.Thr813Profs) | 14.07 | 1/13103 | 25 | 179 | 1/13 | 1/21 | 1/10 | 1/23 |
| Lipoprotein lipase deficiency | LPL:c.701C>T (p.Pro234Leu) | 14.27 | 1/12553 | 6 | 99 | 1/50 | 1/47 | 1/67 | 1/40 |

\*: Autosomal recessive spastic ataxia of Charlevoix-Saguenay
\*\* : Leigh syndrome, French-Canadian type
\*\*\*: Agenesis of corpus callosum and peripheral neuropathy

**Table 2.** New founder pathogenic variant causing primary ciliary dyskinesia.

Autosomal recessive disorders
| Variant | TMRCA (generations) | GnomAD NFE | CARTaGENE |  |  | ISGen |  |
| --- | --- | --- | --- | --- | --- | --- | --- |
|  |  |  | WGS | ARG imputation | TOPMED imputation | Bas Saint Laurent | Côte du Sud |
| HYDIN: c.10426C>T p.(Arg3476Ter) | 12.46 | 1/12810 | 1 | 27 | 31 | 1/303 | 1/301 |

### Carrier frequency in 24 Quebec historical regions

To expand our frequency estimation to other regions of Quebec where sampling within CARTaGENE was not as complete, we took advantage of the availability of the BALSAC genealogy and the presence of 7.894 CARTaGENE individuals in this pedigree. We linked CARTaGENE carriers to the BALSAC genealogy, and we identified a set of probable carrier founders by climbing the pedigree to find common genealogical ancestors using ISGen (9). After identifying candidate founders, we performed allele dropping simulations with ISGen to estimate allele frequencies in 24 historical regions in Quebec (9). In detail, we performed 100k climbing simulations from carriers to founders, and used allele dropping from these 100k trees to estimate allele frequencies [see (9) for details]. We computed confidence intervals using bootstrapping with 100k iterations for individuals within each region.

### Characterizing a rare variant causing primary ciliary dyskinesia in Quebec

The *HYDIN c*.*10426C>T* pathogenic variant is responsible for primary ciliary dyskinesia (PCD – a rare, mostly autosomal recessive condition of motile ciliary dysfunction leading to chronic suppurative respiratory disease) in several cases in Quebec (34). We applied our ARG-imputation method, checking if the pattern within the ARG is consistent with a single founder event, and we estimated the variant age. In addition to CARTaGENE carriers, we linked 7 *HYDIN c*.*10426C>* homozygous patients recruited at the McGill University Health Centre to the BALSAC genealogy. We estimated the frequency across 24 regions of Quebec as described before but added information about these homozygous patients.

## Results

### TMRCAs and global frequencies of putative founder pathogenic variants Testing putative founder pathogenic variants

It has been hypothesized that a few variants in Quebec were introduced by single carriers – these are often referred to as founder pathogenic variants. This hypothesis can be validated in two independent ways: first, all regional carriers should share a recent common ancestor. Second, carrier frequencies in the source populations should be low enough to make the single founder event likely. We, therefore, compared estimated TMRCAs and global allele frequencies for 9 putative founder pathogenic variants in SLSJ. 7 out of 9 cases are consistent with a unique recent common ancestor that settled in Quebec (TMRCA < 17 generations, Table 1 and Supplementary Figure S1, GNOMAD Non-Finnish European (NFE) frequencies less than 1/5000). Here each generation is 29 years, as described in (35). The other two cases, *FAH*: c.1062+5G>A and *GNPTAB*: c.3503_3504delTC have older TMRCAs of 26 and 31 and showed higher GnomAD NFE frequency (≃ 1/1000). Both are thus inconsistent with a single introduction. Further, variants with TMRCA < 17 generations are only present in the NFE population in GnomAD. In contrast, variants *FAH*: c.1062+5G>A and *GNPTAB*: c.3503_3504delTC are present in Finnish (Europeans) and South Asians in GnomAD. The ARG-based TMRCAs do not use information about global gnomAD allele frequencies. The two analyses provide independent and consistent indication about which variants likely resulted from single introductions.

### ARG-imputation

After validating the TMRCA estimates, we focused on the ARG’s topology. We measured clade mixing scores to determine if ARG clades separated carriers from non-carriers. In 7/9 pathogenic variants, these clades only contain true carriers (Clade mixing = 1, Supplementary Table S1). We leveraged these high-quality clades to perform genotype imputation of nine putative founder pathogenic variants in SLSJ (Table 1). We then compared ARG-imputation against imputation by Minimac4 (Supplementary Figure S2). We detected 1,245 concordant positives and 49 discordant positives (imputed by the ARG and not by the reference panel). Moreover, 311 individuals were discordant negatives (imputed by Minimac4 and not the ARG) and 27,728 were classified as negative for both strategies. In addition, we compared the posterior probabilities of carriers imputed by Minimac4, demonstrating that most of them have high posterior probabilities (0.8-1, Supplementary Figure S3). Finally, Kappa agreement was 86.73%, and accuracy was 98.77%, indicating a high agreement between strategies (Supplementary Figure S4).

To further validate our imputation method, we experimentally genotyped imputed carriers of two variants: *PMS2 c*.*2117del* and *ATM c*.*7388_7389insAlu p*.*Leu2463fs*, with 22 and 21 imputed carriers, respectively. For *PMS2 c*.*2117del*, direct genotyping identified 21 carriers. ARG-imputation identified 21 true positives and one false positive (Supplementary Figures S2). In addition, agreement metrics such as precision (positive predictive value), recall (sensitivity), specificity, negative predictive values, and Kappa show a good performance of ARG-imputation to distinguish carriers and non-carriers (Supplementary Figure S4). On the other hand, Minimac4 identified 20 true positives, one false positive and one false negative (Supplementary Figure S2). ARG-imputation outperformed Minimac4 for the *PMS2 c*.*2117del* variant.

For *ATM c*.*7388_7389insAlu p*.*Leu2463fs* pathogenic variant, direct genotyping detected 21 carriers. ARG-imputation identified 21 true positives and 3993 true negatives. The *ATM c*.*7388_7389insAlu p*.*Leu2463fs* pathogenic variant was not detected by high-depth WGS with the DRAGEN pipeline, demonstrating the flexibility of ARG-imputation to incorporate carrier status from various sources, rather than relying solely on WGS reference panels.

### Genotype-phenotype association analysis with *LPL: c*.*701C>T* carriers

After validating our ARG-imputation approach, we aimed to evaluate whether these imputed rare variants were sufficient to capture new associations. We focused on the *LPL* gene for three reasons: the existence of a variant *c*.*701C>T* enriched in QFC, the observation that other variants in that gene display mild phenotype in the heterozygous state, and the availability of blood measurement data in CARTaGENE (29,36). We found a positive association between triglycerides (Beta: 1.26, 95% C.I: 1.02-1.49) and imputed allele dosage (Figure 2). We identified a negative association with HDL-C levels (Beta: -0.28, 95% C.I: -0.35 to -0.20). We did not find a significant association with LDL-C levels (p-value: 1). These findings indicate that *LPL c*.*701C>T* disrupts lipid transport and metabolism involving triglycerides and HDL-C as a heterozygote state, similar to other pathogenic *LPL* variants.

**Figure 2.**
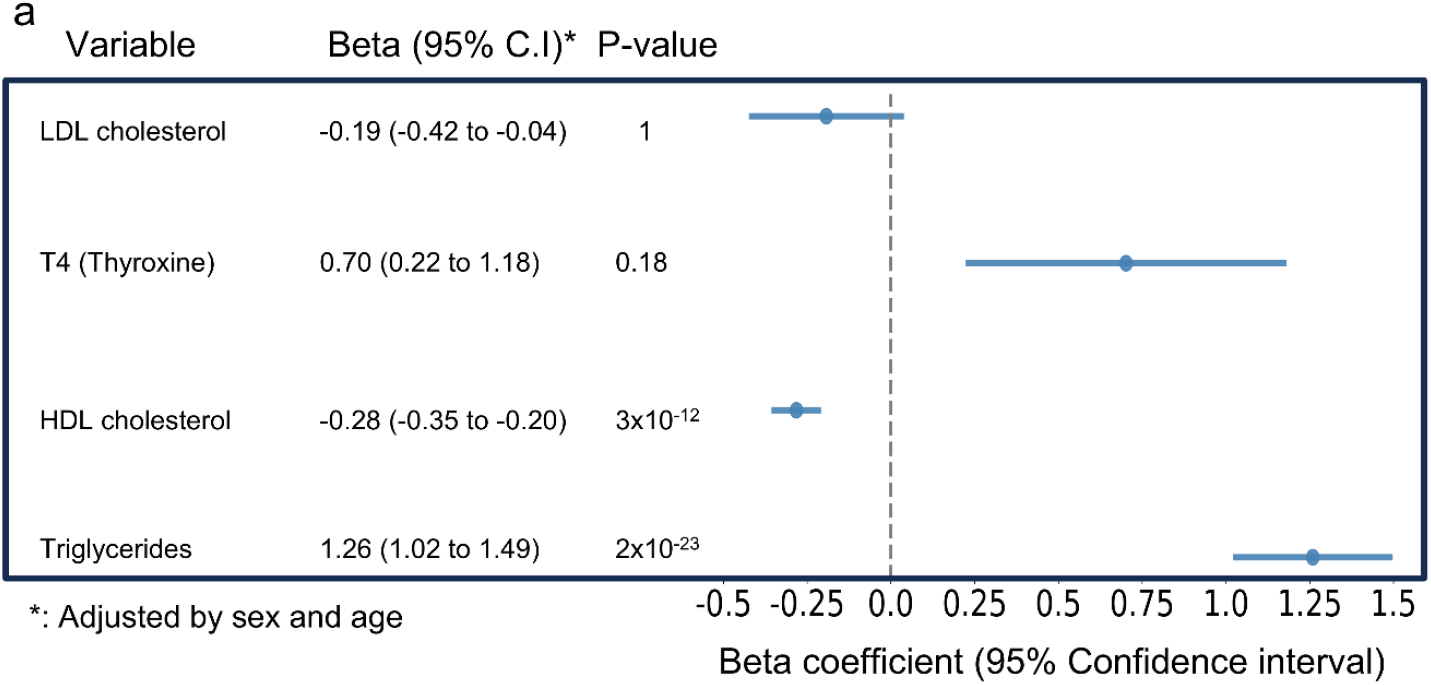
Association between *LPL c*.*701C>T* variant and blood measurements. Phenotype association analysis shows significantly higher levels of serum Triglycerides, T4, and lower levels of HDL-C (b) for individuals with a higher posterior probability of being carriers of *LPL c*.*701C>T* pathogenic variant. Linear regression included sex and age as covariates. P-value were adjusted by Bonferroni correction assuming independent tests.

### Carrier frequency in SLSJ

We used imputed data to estimate carrier frequency in SLSJ. 8/9 founder mutations in Saguenay show similar carrier rates in Saguenay between WGS, ARG-imputation, ISGen, and literature reports (pairwise Fisher exact test, Supplementary Table S2). The *LRPPRC:c*.*1061C>T* variant showed a significant difference between WGS and ARG-imputation (1/125 vs 1/33, P-value= 0.031). Additionally, as expected, these variants have a higher frequency in SLSJ, Charlevoix, and Côte Nord (Supplementary Figure S5) regions relative to the rest of Quebec, due to shared ancestral founder events in the Charlevoix region.

### Founder pathogenic variant causing PCD in the Quebec population

We recently identified a new potential founder pathogenic variant in Quebec causing PCD (*HYDIN c*.*10426C>T)*. The estimated TMRCA for *HYDIN c*.*10426C>T* was 12.4 generations, corresponding to 361 years ago, after the arrival of French individuals in Quebec. Regional frequency estimation using ISGen identified two regions with higher carrier frequency: Bas-Saint-Laurent (1/303) and Côte-Du-Sud (1/301), which is located mainly in the neighbour region of Chaudière-Appalaches but also covers a part of Bas-Saint-Laurent, with a descending gradient in nearby areas (Figure 3b).

**Figure 3.**
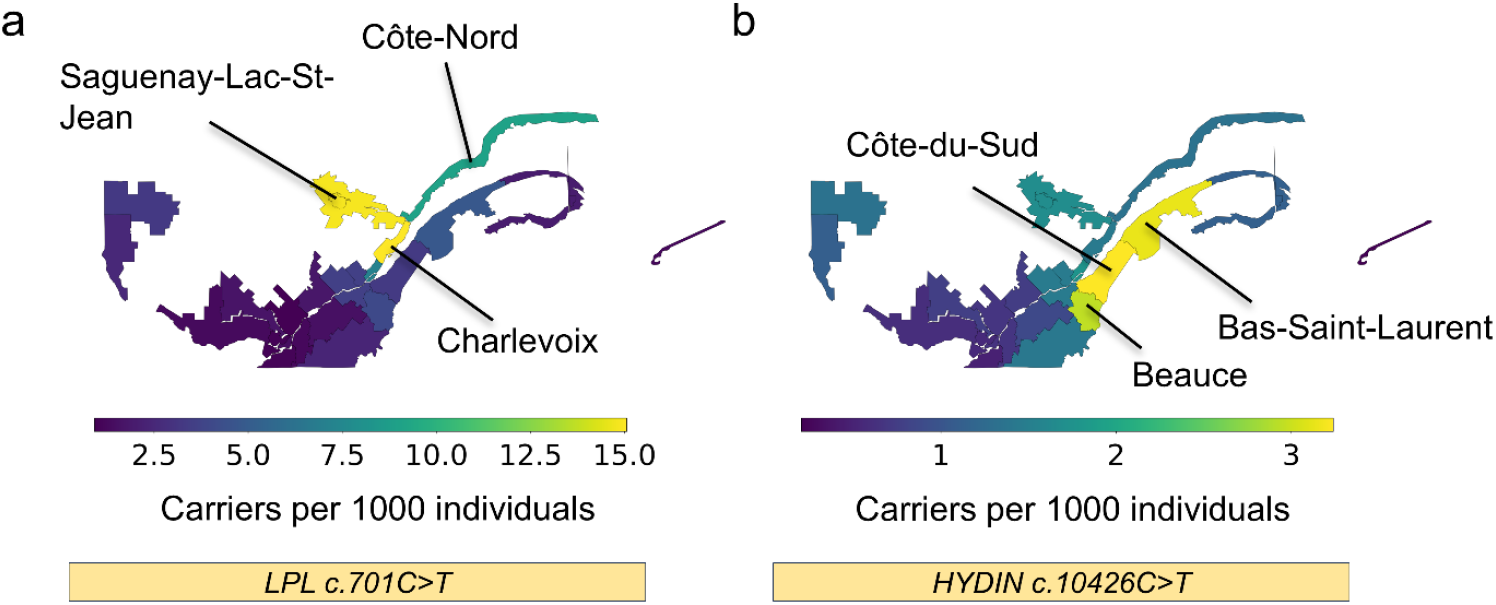
Regional carrier frequency estimation with ISGen. Quebec heatmap with 24 historical regions, scale corresponds to number of carriers per 1000 individuals. a) Carrier frequency of *LPL c*.*701C>T*, showing a higher carrier rate in SLSJ, Charlevoix and Côte-Nord. b) Carrier frequency distribution for *HYDIN c*.*10426C>T* pathogenic variant, showing a higher rate in Bas-Saint-Laurent and Côte-du-Sud.

## Discussion

Founder populations present unique opportunities to study the emergence and propagation of genetic variation. Here, we argue that the ancestral recombination graph provided an excellent tool to combine genotype-inferred haplotype structure with sequencing information to characterize rare variant function and evolution in such populations.

The accuracy of the ARG inferred by ARG-needle has not been widely tested (10). In particular, the accuracy of time estimates for internal nodes representing common ancestors is uncertain. We assessed the consistency of the inferred TMRCAs with the ARG in nine known founder pathogenic variant carriers in SLSJ. TMRCA estimates are consistent with allele frequencies in GnomAD for non-Finnish Europeans (MAF<1/5000 suggests a single introduction into the Quebec population), and with the timing of the known founder events in Quebec.

This independent validation suggests that ARG-needle timing estimates for haplotypes shared by many individuals in the recent past are reasonable. Concretely, we can use the ARG to rapidly identify variants resulting from single introductions. In the case of multiple introductions, subclades in the ARG with deep MRCAs may be identified as likely independent introductions.

We further found agreement between ARG clades inferred from genotype data and clades inferred from the carrier status of rare variants (Table 1). We, therefore, used the ARG to perform genotype imputation.

Our ARG-imputation method is very similar to the one implemented by Zhang et al (10). Both consider branch lengths and the ARG’s topology to give a weighted average of implied genotypes (comparable to our posterior probability). For rare variants, ARG-imputation showed a higher performance than standard imputation methods (IMPUTE4), as demonstrated by Zhang et al (10). Our results also showed high performance for ultra-rare variants, highlighting the importance of ARG-imputation for low-frequency variants, especially in founder populations. We were able to easily perform imputation for a variant that was unavailable from a WGS reference panel, using a small number of directly-genotypes cases.

In some cases, individuals who were identified as non-carriers by WGS data would have been imputed as carriers by our method (i.e., they coalesce with true carriers below the MRCA in the ARG). Four events could explain this scenario: 1) an error in the ARG inference, 2) back mutation or reversion, 3) and gene conversion 4) sequencing error. A deeper analysis of mismatches is a promising avenue to study the recent history of rare variants.

After validating our ARG-imputation approach, we evaluated the association between lipid measurements and *LPL c*.*701C>T* carriers. Carriers indeed showed a significant increase in triglycerides, decreased HDL levels in serum, and a higher risk of developing metabolic syndrome (Figure 2).

The largest study on *LPL* loss of function (LOF) carriers and lipid levels found effects comparable to the *LPL c*.*701C>T* variant reported here, with a positive association between carrier status and triglycerides levels (beta= 1.086 mmol/L, p-value= 0.01) and a negative association with HDL-C (beta= -0.19 mmol/L, p-value=0.001). This demonstrates that the effect size for *LPL c*.*701C>T* is comparable to other *LPL* pathogenic variants (37). Due to the low frequency of this variant outside Quebec, this is the first association study result in *LPL c*.*701C>T* carriers.

Finally, we leveraged imputed carriers to perform pedigree-based regional frequency estimation. The ARG provides information about whether a variant was introduced once or multiple times. This is essential for regional frequency estimation using ISGen, which assumes a single founder introduction (9). We tested the accuracy of regional frequency estimation by studying nine pathogenic variants for which regional fand provincial frequencies are already known (Table 1). Patterns given by ISGen (Figure 3 and Supplementary Figure S5) agreed with literature reports that show higher frequency in Saguenay, Charlevoix, and Côte-Nord (21,38–45). Because it imputes carrier status in both sampled and unsampled individuals, ISGEN can partially overcome sampling biases(9). Literature reports of carrier frequencies vary in their fine-scale sampling biases. The proposed pipeline offers an opportunity to extrapolate carrier frequencies outside the sampling area, which may lead to more accurate genetic counseling and screening in eligible populations.

In conclusion, we argue that ARGs should become part of the standard toolkit for the study of rare variants, especially in founder populations, whether or not genealogical data is available.

## Data and code availability

Scripts to perform ARG-imputation are available here: https://github.com/almejiaga/ARG-imputation. For ISGen regional frequency estimates, a fork repository is available: https://github.com/almejiaga/ISGen. The CARTaGENE genetic and phenotypic data is available from https://cartagene.qc.ca/en/researchers/access-request.html after scientific and ethical review. The CARTaGENE allele frequency information is publicly available from: https://cartagene-bravo.cerc-genomic-medicine.ca/.

## Acknowledgments

The authors would like to thank all the patients and their families at the McGill University Health Center for their involvement in identifying a novel *HYDIN* pathogenic variant.

## Author contributions

A.M-G. performed the ARG imputation, allele frequency analysis and genotype-phenotype associations. S.G. designed and supervised the work. A.M-G. and S.G. adapted the ARG-based imputation method from ARG-needle. A.D.P. performed the UMAP-based clustering to define the SLSJ genetic group. G.S., D.D., L.B. and A-J.S. contributed to identifying the *HYDIN* variants and to the conceptualization of this study. W.F., N.H., A.L.C. and G.C. performed the experimental validation for PMS2 and ATM variants. K.S.L. and G.L. performed functional annotation of the putative founder variants and pre-processed genetic data from CARTaGENE. V.C. and D.T. performed imputation with the Quebec reference panel. S.G. and A.M-G. wrote the initial manuscript. All authors edited and reviewed the final version of the manuscript.

