## Supplemental_information for "Using the ancestral recombination graph to study the history of rare variants in founder populations"

### Supplemental Figures

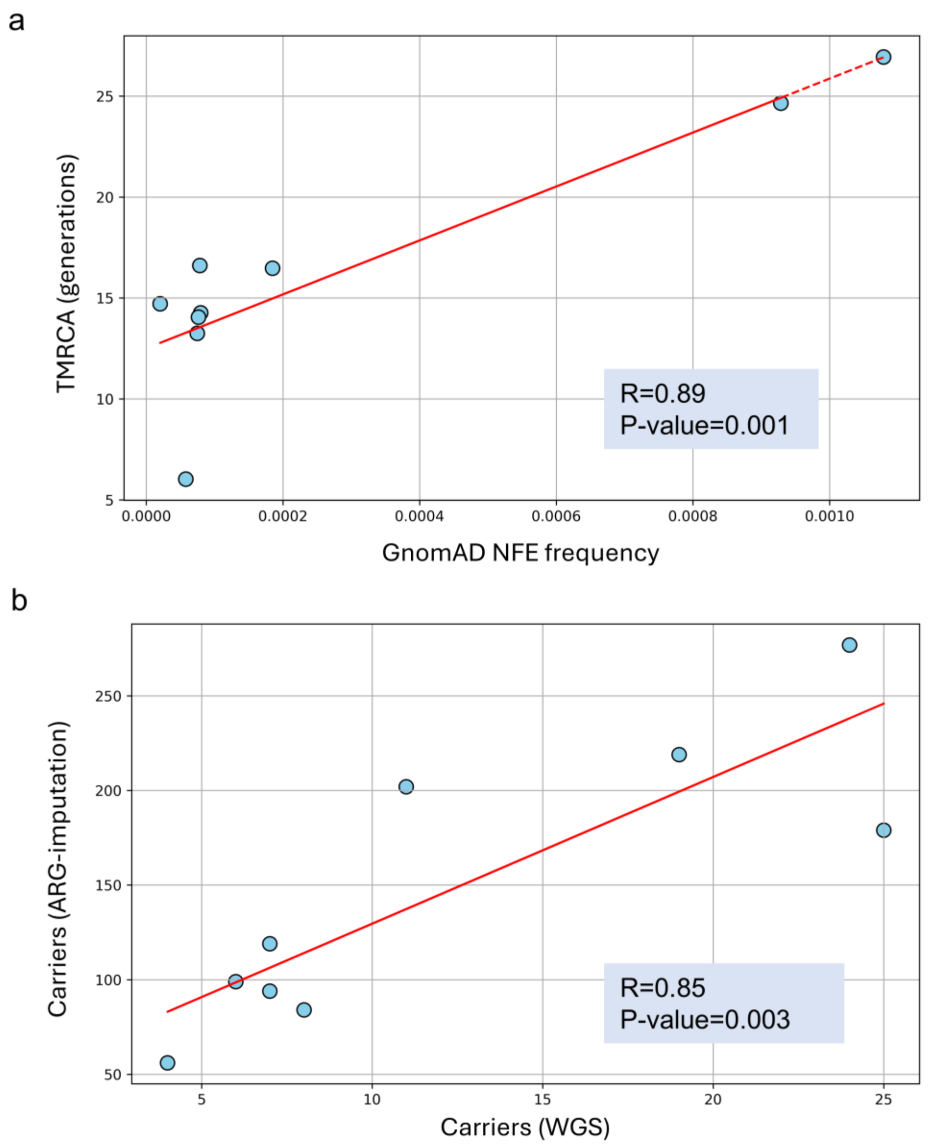


Figure S1: Positive correlation between frequency for non-Finnish Europeans in GnomAD and TMRCA, and the number of carriers in WGS and ARG-imputation. a) scatter plot between TMRCA and frequency in GnomAD for non-Finnish Europeans for 9 putative founder variants in SLSJ, showing a high correlation (R=0.89). A Linear relationship between the number of carriers in the CARTaGENE cohort by WGS and after ARG-imputation for 9 putative founder variants in SLSJ.

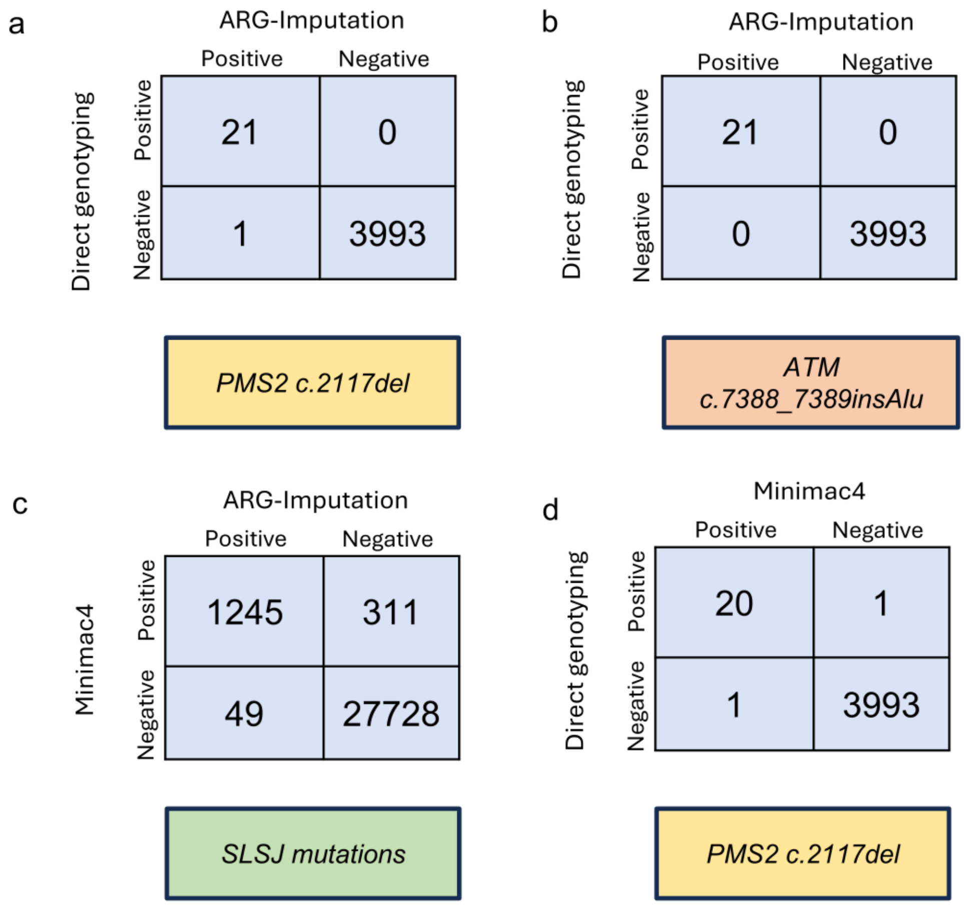


Figure S2. ARG-imputation validation with direct genotyping and TOPMED imputation. Contingency tables comparing ARG-imputation against direct genotyping (gold standard) for two mutations (a and b) and 9 putative founder mutations in SLSJ. a) PMS2 c.2117del mutation. ARG-imputation captures 21 true positives, 1 false positive and 3993 true negatives. B) ATM c.7388_7389insAlu. ARG-imputation captures 21 true positives and 3993 true negatives. c) 9 putative founder mutations in SLSJ. We identified 1245 concordant positives, 49 discordant positives, 311 discordant negatives, and 27728 concordant negatives. d) PMS2 c.2117del mutation imputation with Minimac4 compared to direct genotyping (gold standard). Minimac4 captures 20 true positives, 1 false positive and one false negative.

a

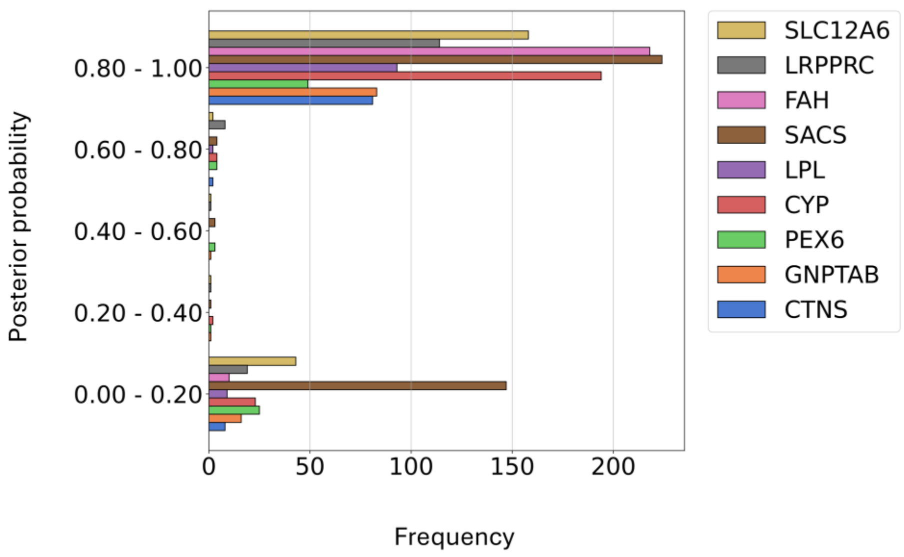


Figure S3. Comparison of imputation with ARG and Minimac4 using a Quebec reference panel. Posterior probabilities of individuals inferred as carriers for 9 putative founder mutations using Minimac4 are displayed. ARG-imputation is concordant with Minimac4 (posterior probability between 0.80 - 1). In a few cases, ARG-imputation does not capture Minimac4 carriers (posterior probability between 0 – 0.20).

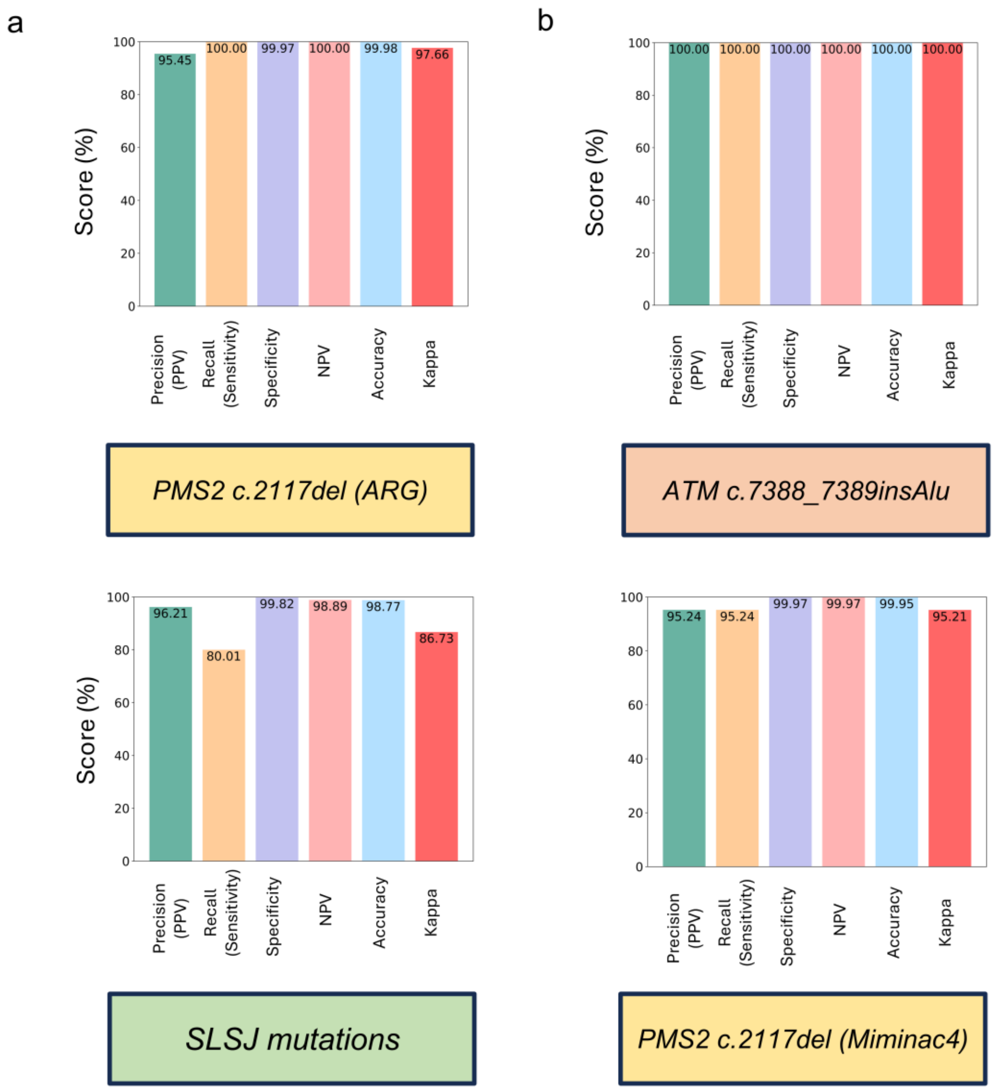


Figure S4. Performance of ARG-imputation compared to direct Genotyping and Minimac4 imputation using a Quebec reference panel. a) PMS2 c.2117del mutation shows a high precision, sensitivity, and specificity, among other concordance metrics. b) ATM c.7388_7389insAlu mutation demonstrated a 100% performance in all metrics. c) Comparison between imputation using the ARG and Minimac4 with a Quebec reference panel, using Minimac4 as a gold standard. Kappa and Recall showed 86% and 80% respectively. The rest of the metrics are above 95%. d) PMS2 c.2117del imputation by Minimac4 shows lower performance than ARG-imputation in terms of PPV, recall, specificity, NPV, accuracy and Kappa agreement.

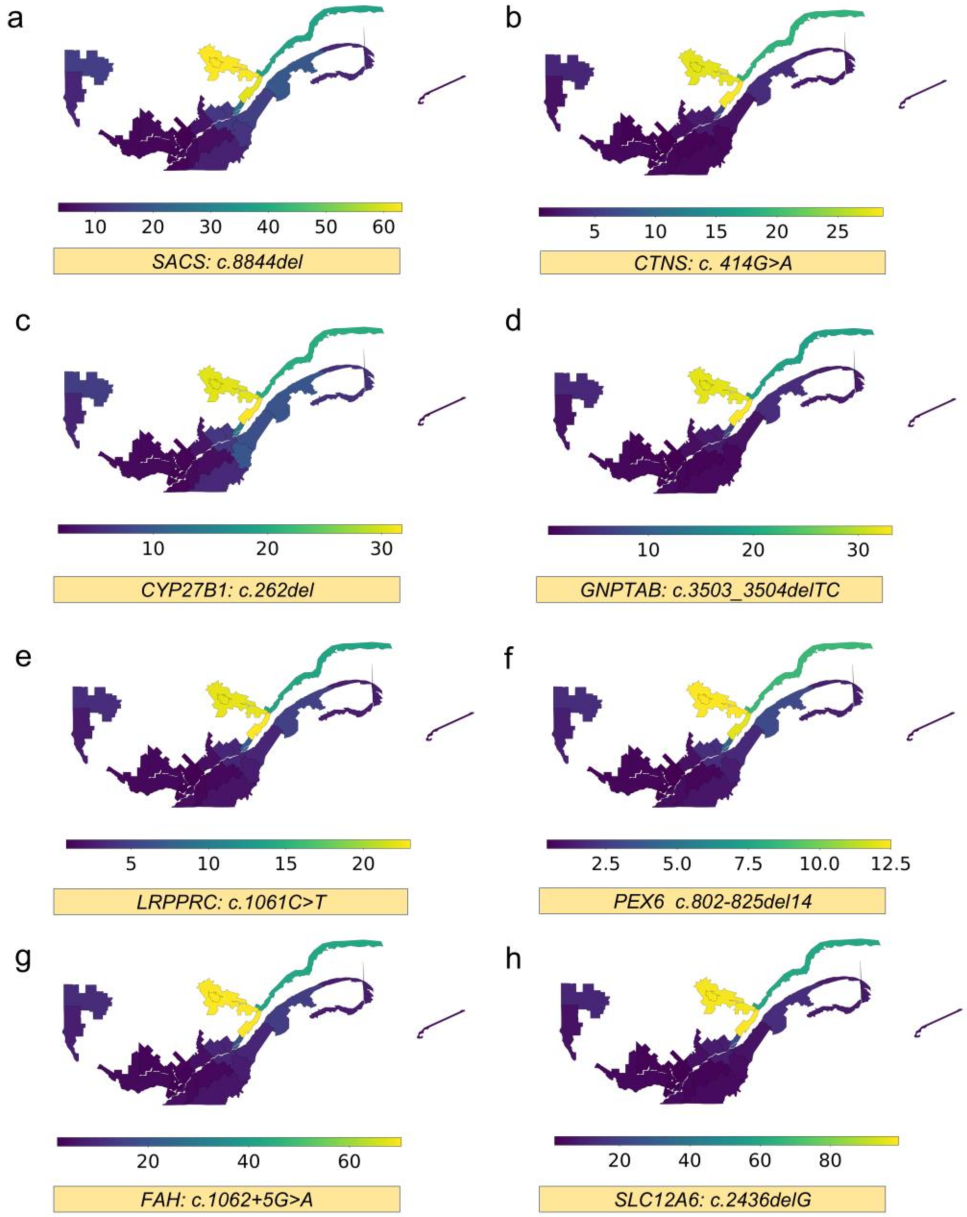


Figure S5. Regional carrier frequencies for 8 putative founder mutations in SLSJ. Heatmap with the number of carriers per 1000 individuals in 24 historical regions of Quebec (as defined in the BALSAC project). All variants have the same pattern: higher frequency is SLSJ, Charlevoix and Côte-Nord.

Supplemental Tables

Table S1:

| Table S1: Imputation score, leave one out and Jackknife bias estimation for 9 putative founder mutations in SLSJ | | | |
|---|---|---|---|
| Mutation | Imputation Q score | LOO RMSE | Jackknife bias |
| GNPTAB: c.3503_3504delTC | 1 – 0/8 | 0.0419 | -0.1298 |
| PEX6 c.802-825del14 | 1- 0/4 | 0.114 | -1.18 |
| FAH:c.1062+5G>A | 1 – 0/19 | 0 | 0 |
| CTNS:c. 414G>A (p.Trp138X) | 1 – 3/7 | 0 | 0 |
| SACS | 1 – 2/24 | 0.0889 | -0.203 |
| LRPPRC:c.1061C>T (p.Ala354Val) | 1 – 0/7 | 0 | -2.84e-14 |
| (CYP27B1):c.262del (p.Val88fs) | 1 – 1/11 | 0.0862 | -1.1408 |
| SLC12A6:c.2436delG (p.Thr813Profs) | 1- 0/25 | 0.00270 | -0.0226 |
| LPL:c.701C>T (p.Pro234Leu | 1 – 0/6 | 0.0826 | -1.1408 |

Table S2:

| Table S1: P values Fixer exact test Saguenay Carrier frequency | | | | | | |
|---|---|---|---|---|---|---|
| **Mutation** | **WGS vs ARG-imputation** | **WGS vs Isgen** | **WGS vs Literature** | **ARG-imputation vs ISGen** | **ARG-imputation vs Literature** | **ISGen vs Literature** |
| GNPTAB: c.3503_3504delTC | 0.47 | 0.56 | 1.00 | 0.46 | 0.54 | 1.00 |

| | | | | | | |
|---|---|---|---|---|---|---|
| PEX6 c.802-825del14 | 0.52 | 1.00 | 1.00 | 1.00 | 1.00 | 1.00 |
| FAH:c.1062+5G>A | 1.00 | 0.70 | 0.14 | 0.71 | 0.17 | 0.44 |
| CTNS:c. 414G>A (p.Trp138X) | 0.82 | 1.00 | 1.00 | 1.00 | 0.58 | 1.00 |
| (CYP27B1):c.262del (p.Val88fs) | 1.00 | 0.60 | 1.00 | 0.62 | 1.00 | 1.00 |
| SACS:c.8844del (p.Ile2949fs) | 0.19 | 1.00 | 1.00 | 1.00 | 0.72 | 1.00 |
| LRPPRC:c.1061C>T (p.Ala354Val) | 0.031 | 1.00 | 0.35 | 1.00 | 1.00 | 1.00 |
| SLC12A6:c.2436delG (p.Thr813Profs) | 0.04 | 0.72 | 0.49 | 0.18 | 1.00 | 0.31 |
| LPL:c.701C>T (p.Pro234Leu) | 1.00 | 1.00 | 1.00 | 1.00 | 1.00 | 1.00 |

### Supplemental Methods

**Leave one out experiment for imputation robustness**
To assess the robustness of ARG-imputation to the number of carriers used for defining the MRCA and the posterior probability, we implemented a leave-one-out approach. For a variant x with carriers forming the set, we formed subsets of these carriers, leaving one individual out in each of them. After, we applied the imputation approach as explained before. To measure accuracy, we compared the posterior probabilities assigned using the whole set of carriers against each leave-one-out subset. We then estimated the root mean squared (RMSE) error for every subset and selected the highest one.

**Jackknife bias for ARG-imputation**
To assess the bias in the ARG-imputation, we followed a jackknife sampling approach on the carriers used for imputation. We then formed the same C subsets leaving one individual out and then performed ARG-imputation. After, we summed the posterior probabilities of the individuals. Finally. We estimated the mean of this sum and use it as the jackknife parameter to estimate the bias, as described in (1).

**Comparing carrier frequencies between WGS, ARG-imputation, ISGen and literature**
We employed a Fisher exact test to compare the carrier frequency estimated by WGS, ARG-imputation, ISGen and literature. However, when comparing WGS and ARG-imputation, these two estimations are not independent. We then subtracted all the WGS individuals from the ARG-imputation estimates to meet the independence assumption in the Fisher exact test.

We found an apparent discrepancy in the carrier frequency for the *LRPPRC:c.1061C>T* mutation between WGS and ARG-imputation. However, only two CARTaGENE WGS carriers are from the genetically inferred SLSJ status. While the difference is statistically significant, subtle differences in regional sampling may explain the difference. Evaluating the statistical significance of this difference is challenging because the same sequence data used to learn the carrier status was used for imputation. In addition, the same genetic data used for imputation was used to assign SLSJ status with a genetic clustering algorithm (2). Thus, statistical independence of a small number of observations is hard to establish. Further, distinguishing between individuals from SLSJ and Charlevoix is challenging due to migration history, where most of the founders of SLSJ came from Charlevoix and, therefore, share common ancestors. With these caveats in mind, ARG-imputation captures many more carriers in the whole dataset, providing a more detailed estimate of carrier rates (119 carriers).
